# Centennial clonal stability of asexual *Daphnia* in Greenland lakes despite climate variability

**DOI:** 10.1101/2020.07.22.208553

**Authors:** Maison Dane, N. John Anderson, Christopher L. Osburn, John K. Colbourne, Dagmar Frisch

## Abstract

Climate and environmental condition drive biodiversity at many levels of biological organisation, from populations to ecosystems. Combined with palaeoecological reconstructions, palaeogenetic information on resident populations provides novel insights into evolutionary trajectories and genetic diversity driven by environmental variability. While temporal observations of changing genetic structure are often made of sexual populations, little is known about how environmental change affects the long-term fate of asexual lineages. Here, we provide information on obligately asexual, triploid *Daphnia* populations from three Arctic lakes in West Greenland through the past 200-300 years to test the impact of a changing environment on the temporal and spatial population genetic structure. The contrasting ecological state of the lakes, specifically regarding salinity and habitat structure may explain the observed lake-specific clonal composition over time. Palaeolimnological reconstructions show considerable environmental fluctuations since 1700 (the end of the Little Ice Age), but the population genetic structure in two lakes was almost unchanged with at most two clones per time period. Their local populations were strongly dominated by a single clone that has persisted for 250-300 years. We discuss three possible explanations for the apparent population genetic stability: (1) the persistent clones are general purpose genotypes that thrive under broad environmental conditions, (2) clonal lineages evolved subtle genotypic differences that are unresolved by microsatellite markers, or (3) epigenetic modifications allow for clonal adaptation to changing environmental conditions. Our results will motivate research into the mechanisms of adaptation in these populations, as well as their evolutionary fate in the light of accelerating climate change in the polar regions.

## Introduction

Climate and the environment are major drivers of biodiversity in the broad sense. In addition to higher levels of biological organisation (e.g. communities, ecosystems), these environmental drivers affect species at the population level, including their temporal and spatial genetic structure and diversity [1,2]. The responses of population genetic structure and diversity to environmental forcing need to be considered at a range of temporal and spatial scales to fully evaluate how global change processes affect species’ distributions and adaptive capacities [3,4]. Long-term, field-based perspectives that span pre- and post-disturbance time periods (typically > 100 years to cover pre-industrial times) are required to balance experimental and laboratory approaches at elucidating ecological responses and genetic adaptation to climate change and chemical pollutants [5]. Studies that combine palaeoecological data with molecular data are especially promising to advance our understanding of past and current adaptation to environmental shifts [3,6].

Palaeolimnological approaches have a long history of providing detailed reconstructions of ecosystem variability at a range of temporal scales (10^0^ to 10^3^ yrs) and across trophic levels [7]. Recent methodological progress combined with a better understanding of preservational constraints and artefacts (e.g. seeds, eggs, cysts) has allowed the use of bulk (environmental) and compound-specific (organismal) DNA to address questions about long-term genetic trends, variability and adaptation in aquatic ecosystems [5,8]. The application of DNA extracted from dormant propagules can also be coupled with resurrection approaches [8,9]. In optimal conditions, the palaeogenetic record allows the reconstruction of temporal patterns of genetic structure and diversity of aquatic populations and communities. These palaeogenetic reconstructions can be coupled with more standard palaeoecological proxies (microfossils, geochemical markers, etc.) that reflect environmental and climatic drivers and forcing.

Genetic diversity is often reduced during times of environmental disturbance following strong selection and resulting population bottlenecks [10]. Shifts in genetic structure associated with rapid environmental change have also been observed in long-term studies [11,12]. Likewise, the spatial genetic structure of populations is strongly related to environmental gradients, creating patterns of isolation by environment [13]. Many species have undergone adaptive evolution to shifts in environmental conditions [14]. While these observations are generally made in sexual populations, little is known about long-term dynamics in the genetic make-up of asexual populations. Under an environmental change scenario, asexual lineages may be at a disadvantage compared with sexuals, by shuffling their genetic material thus creating the variation needed to fuel selection [15,16]. However, the potential of asexual lineages to persist during the rapid change currently recorded across ecosystems remains unclear. Obligate parthenogenesis in the *Daphnia pulex* species complex involves the production of apomictic eggs [17,18] that are deposited in an ephippium. This chitinous structure is derived from maternal tissue, and each contains two dormant eggs, which in asexual populations are genetically identical. Ephippia with unhatched eggs can be extracted from the sediment and their DNA examined, allowing for a detailed study of temporal population genetic structure of *Daphnia* populations [12,19–22]. Long-term studies at the scale of centuries on the fate of asexual *Daphnia* lineages do not exist. However, various asexual *Daphnia* lineages have been recorded for 20-30 years in the Canadian Subarctic [23], and for over 70 years in an African lake [20]. Contrasting the idea of centennial persistence of asexual lineages is the age estimate of only decades [24] for asexual lineages of the North American *Daphnia pulex-*complex that arose by contagious asexuality [25].

Arctic lakes in Greenland provide an ideal system to study long-term dynamics of asexual populations. Many lakes are fishless and inhabited by populations of the keystone herbivore *Daphnia*, in particular the large-bodied *Daphnia pulex*-complex [7]. Arctic *Daphnia* populations are generally obligate asexuals [26,27], and many of these lineages are triploids [28]. In the Illulissat area of West Greenland, clonal richness of *Daphnia pulex* populations was between 4 and 5 genotypes [29,30]. The lakes along Kangerlussuaq in SW Greenland are particularly well-studied in relationship to recent climate change, with evidence for a dramatic increase in environmental forcing of these sensitive habitats [31]. In the context of the present study, the Kangerlussuaq area is a natural experimental site, in that direct anthropogenic impacts are limited or absent. But there have been substantial environmental changes reflecting climate (temperature, precipitation and altered seasonality) as well as related landscape changes including soil erosion, dust deflation, lake shrinkage, and terrestrial vegetation changes [32]. Kangerlussuaq, in contrast to most Arctic regions, was cooling throughout much of the 20^th^ century. However, since 1996 the area has undergone pronounced climate change, including rapid warming (> 2 °C) over the last 15 years, partly in relation to changes in the Greenland Blocking Index [31].

In order to advance our understanding of the fate of asexual populations in the light of current environmental change at high latitudes, there is a need for empirical data, especially on decadal to centennial timescales. Here, we examine genetic diversity in polymorphic microsatellite DNA, and spatio-temporal genetic structure of three *Daphnia* lake populations in West Greenland near Kangerlussuaq and their relationship with the known regional environmental history. We test the hypothesis that rapid environmental change over the last 200-300 years has driven change in temporal and spatial population genetic structure of asexual *Daphnia* populations at the individual lake and landscape scale.

## Material and Methods

### Study area

The Kangerlussuaq area of SW Greenland is a major lake district with thousands of lakes, with a reasonably well understood regional limnology [33]. This, coupled with preliminary zooplankton surveys and the fact that a number of the lakes have been the subject of palaeolimnological analyses (e.g. [34,35]) meant that sites could be chosen to optimize ephippia recovery, namely deep seasonally anoxic basins. Braya Sø (SS4) was also known to have high concentrations of sediment biomarkers [36]. The area around the head of the fjord (Fig. 1) has low effective precipitation and is characterized by limited hydrological connectivity resulting in a number of closed-basin oligosaline lakes (conductivity range 1000–4000 μS cm^-1^). In general, the Kangerlussuaq freshwater lakes at the head of the fjord have conductivities around 200-500 μS cm^-1^ and DOC concentrations ca. 30-50 mg L^-1^; DOC in the oligosaline lakes can be much higher (80-100 mg L^-1^ [37]; nutrient and major ion water chemistry is given in Table S1). The study lakes are located within a few km of each other (Fig. 1). SS1381 and SS1590 are small (21.5 and 24.6 ha respectively) scour basins with maximum depths around 18 m. Both lakes stratify strongly with anoxic hypolimnia. Braya Sø is a larger, meromictic oligosaline lake (73 ha). The ice-free period is from late-May / early-June to late September. The lakes can be considered pristine in comparison to temperate systems in North America and NW Europe and have no direct cultural impact apart from low levels of atmospheric pollution [38]. The catchment vegetation of all lakes is dwarf shrub tundra.

**Fig. 1.**
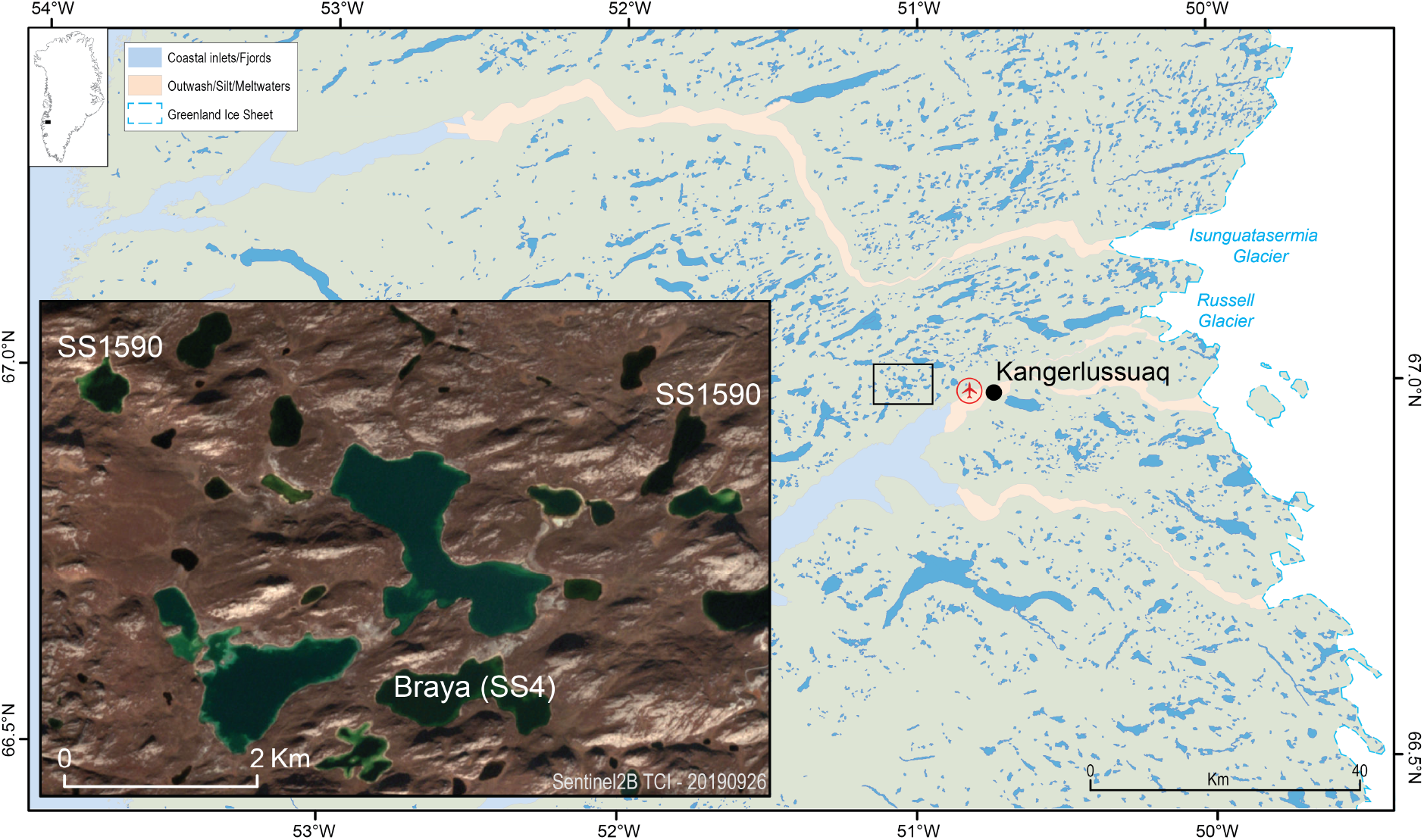
Overview of the study area. Insert shows location of the study lakes SS4 (Braya Sø), SS1590,and SS1381.

The study lakes are presently fishless, which is required for *Daphnia* populations to persist [39]. All the lakes have variable water levels both at inter-annual and decadal timescales, resulting from their sensitivity to the seasonality of precipitation, the extent of the spring melt input and intense evaporation during the summer [40]. Although the lakes are isolated basins today, SS4 was once part of a group of lakes that formed a large palaeolake in the early Holocene [41]; SS1590 would have drained into this larger lake for a brief period. SS1381 is located 6 km to the west of SS1590 and above the level of the highest shoreline of the palaeolake.

West Greenland has undergone considerable environmental change (over different timescales) since deglaciation. Research has focussed on a range of biophysical aspects of the ice-land-water continuum, including glacier mass balance, terrestrial vegetation and herbivore dynamics, and aquatic biogeochemistry [32]. Lake sediment records have provided details of ecological response to climate and landscape change over centennial and millennial timescales, including diatom, pigment and palaeoclimatic reconstructions [34,35,42,43]. Because ephippia recovery was greatest at SS4 and this lake has a well-defined environmental history, discussion of microsatellite variability and environmental history is restricted to this site.

### Sediment sampling and analysis

Short sediment cores (4 replicates per lake) were taken in July 2015 with a 9.5 cm diameter Hon-Kajak sediment corer from the deepest part of each lake. Recovery was around 25-cm at each lake and at all sites included an intact sediment water interface (Fig. S1). Cores were sectioned on shore at 1-cm intervals and samples were placed in individual plastic bags. Although sedimentation rates are low in these lakes (~0.3 cm yr^−1^; refs), a 1-cm interval was used to maximise recovery of ephippia; finer interval sampling would have provided too few ephippia. Samples were kept refrigerated after return from the field until further analysis. A sub-sample of each core slice was analysed for dry weight and organic matter content using standard methods (loss-on-ignition at 550 °C). The cores used in this study were not dated radiometrically but previous cores have been dated using ^210^Pb and ^14^C [43,44]. Organic matter profiles are distinctive at each lake (Fig. S2), allowing among-core correlation and thus chronologies to be transferred from the dated cores to those used in this study. This method is illustrated for SS4 where 4 replicate cores were taken in 2015; core OM profiles are clearly repeatable (Fig. S2). Samples used for microsatellite analysis cover the most recent 200–300 years for SS4 and SS1381. The higher sedimentation rate at SS1590 meant that recovery of ephippia with eggs suitable for microsatellite analysis was confined to the most recent ~30 years based on ^210^Pb analysis (NJ Anderson unpublished).

Previously, the changing abundance of purple sulphur bacteria (PSB) at Braya Sø has been inferred from the variable concentration of the carotenoid okenone in the lake sediment (see [34]). Unfortunately, pigments were not measured on the sediment cores used in this study but we have shown a good agreement between sediment porewater fluorescence of the cores sampled in 2015 and okenone abundance from a 2001 core (Osburn & Anderson unpublished). Therefore, in this study we used the second component (C2) of a PARAFAC model based on the fluorescence excitation emission matrices (EEMs) of porewater chromophoric dissolved organic matter (CDOM) as a proxy for changing PSB abundance. Component 2 exhibits a clear peak centred on 280 nm and is a protein-like fluorescence marker associated with bacteria. Porewater CDOM fluorescence was measured on separate Varian Eclipse spectrofluorometers and fluorescence intensity was calibrated into Raman units.

The organic C flux was calculated as in previous studies [45] and at SS4 the ephippia accumulation rate was estimated as the total number of ephippia in a 1-cm sediment slice divided by the linear sediment accumulation rate (yrs cm^-1^).

### Molecular analysis

The studied species was identified as *Daphnia pulex* sensu lato (in the following: *Daphnia piilex*). and clustered with lineages of the Polar *Daphnia pulicaria* clade based on the mitochondrial ND5 [46] (D. Frisch, unpublished data).

Ephippia were removed from 1-cm-sections of sediment and enumerated for each sediment section as described in Frisch et al. [12]. Eggs were removed from the ephippia and used for DNA extraction (using one egg per ephippium). DNA was isolated from individual eggs with the QIAamp Microkit (Qiagen Inc, Valencia, CA, USA) following the protocol for DNA extraction from tissue specified in the manufacturer’s manual. To facilitate DNA extraction, eggs were perforated with a sterile micropipette tip, breaking the membrane and exposing egg contents to the extraction buffers. The total number of DNA isolates was 92 from three lakes (n = 57 (SS4), n = 12 (SS1590), n = 23 (SS1381)).

Microsatellite loci were amplified in single, 12.5 μl multiplex reactions (Type-it PCR kit, Qiagen Inc, Valencia, CA, USA), using an Eppendorf Nexus Thermal Cycler with thermal cycle conditions recommended in the Type-it PCR kit manual. Ten microsatellite primers (Table S2) representing genome-wide loci were used for genotyping; details in [12,47]). Two primers (Dp90, Dp377) failed to amplify in a consistent manner and were therefore excluded from further analysis. Amplified microsatellites were genotyped on an Applied Biosystems 3730 genetic analyser. We used the microsatellite plugin for Geneious 7.0.6 (https://www.geneious.com) for peak calling and binning. Called peaks were visually inspected and manually adjusted when necessary.

All analyses were run in R version 3.6.2 [48] using the R package *poppr* 2.8.3 [49]. For the population genetic analyses, we excluded all isolates with >5% missing information, yielding a set of 59 isolates (SS4: 28 isolates; SS1381: 10 isolates; SS1590: 21 isolates, Table S3). Analyses included the estimation of allelic diversity for each locus, and the population genetic parameters of estimated multilocus genotypes (eMLG) by rarefaction to the smallest population size of 10 in SS1590.

Population genetic structure inferred by the number of clusters of genetically related individuals was analysed using Discriminant Analysis of Principal Components (DAPC) [50].This procedure involves the computation of a Principal Components Analysis (PCA) followed by Discriminant Analysis (DA). The number of PCs to be used in DA was identified by the function xvalDapc() implemented in *poppr*.

## Results

### Environmental variability and ephippial dynamics

Regional climate in South West Greenland was variable from the end of the Little Ice Age (LIA) with alternating warm and cold periods, and the most recent warm period beginning at the end of the 20^th^ century (Fig. 2A). As example for the in-lake temporal variability in the last centuries, we focus on reconstruction of environmental conditions in SS4. Here, both the OC accumulation rate and the abundance of PSB (as fluorescence PARAFAC component C2) varied over the last 300 years suggesting variable lake production (Fig. 2B). *Daphnia pulex* ephippia flux varied from ca. 2 to 900 ephippia m^2^ yr^-1^ in 1700 to post 1950, peaking at 888 around 1900 (Fig. 2C), indicative of considerable fluctuations in population size.

**Fig. 2.**
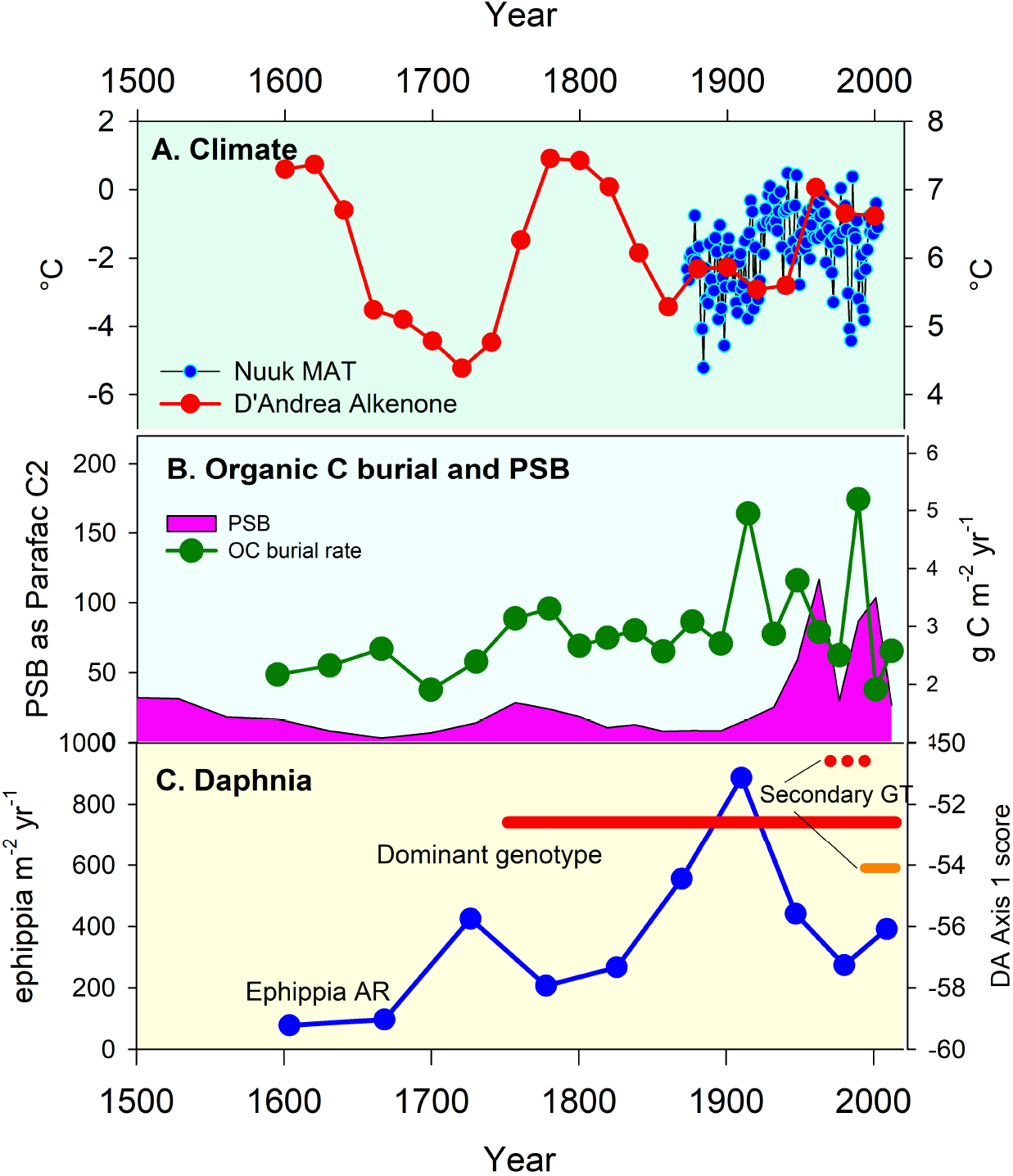
Environmental history at SS4 (Braya Sø) and the response of Daphnia genotypes. A. Climate variability: reconstructed summer lake water temperatures based on alkenones at Braya Sø and Lake E (see [44] for details) and the mean annual air temperature recorded at Nuuk. B. Aquatic production indices for the lake: organic C burial reflects total in-lake production and abundance of purple sulphur bacteria inferred from porewater fluorescence (as Parafac Component C2). C. The stability of the dominant genotype over time period covered by the sediment analyses is indicated by the solid red line; two secondary, minor genotypes (GT) are indicated (solid orange and dotted red lines).

### Population genetic parameters

Genetic diversity and population structure of *Daphnia* populations were studied in three lakes near Kangerlussuaq, SW Greenland (SS4, SS1381, SS1590, Fig. 1), using dormant eggs deposited in the sediment. The covered time period differed between lakes and was 200-300 years in SS4 and SS1381, and ~30 years in SS1590. All *Daphnia* clones were putatively triploid, based on the criterion of three different alleles at a minimum of one locus in each multi-locus genotype (MLG). None of the MLGs included a locus with more than three alleles. Of the eight loci used for genotyping, six (Dp162, Dp291, Dp369, Dp401, Dp437, Dp461) had three different alleles in at least one individual, while two (Dp43, Dp173) had a maximum of two alleles.

Clonal diversity differed between the three lakes (Table 1): of 28 eggs examined in SS4, we found a total of three MLGs (maximum of 2 per time period, Fig. 3A) of which one was dominant throughout three centuries (MLG1). Only a single clone (MLG10) was detected in the 21 isolates from SS1381, which persisted over several centuries, as did MLG1 in SS4. In contrast, the *Daphnia* population in SS1590 was more diverse than the former two lakes with seven MLGs in a total of 10 isolates, of which only one MLG was found during two (non-adjacent) time periods. These differences in clonal diversity and dominance structure are also reflected in the rarefied estimate of clonal diversity (eMLG, Table 1), which in SS1590 was about 3.5 and 6 times higher than in SS4 and SS1381, respectively. No clear temporal pattern was discernible in either of the two lakes with long-term records (Fig. 3A, SS4 and SS1381).

**Fig. 3.**
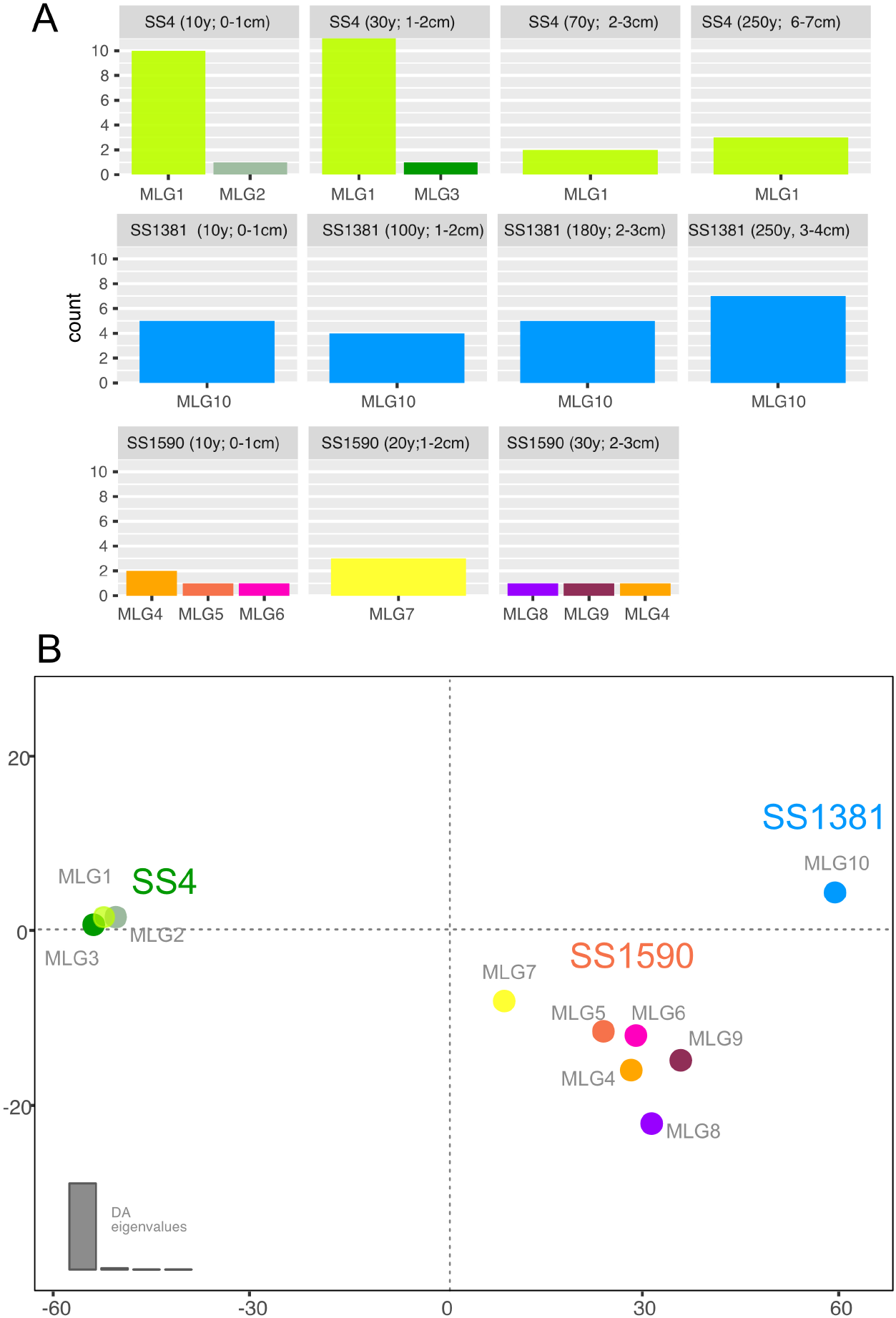
Spatial and temporal population genetic structure of the Daphnia populations in three lakes. To avoid an artificial increase of identical genotypes, only one egg per ephippium was used for microsatellite analysis. A. Abundance of multilocus genotypes (MLG1 to MLG10) with information on lake and sediment age from which eggs were isolated. Each row represents a lake: SS4 (4 time periods), SS1381 (4 time periods), and SS1590 (three time periods). B. DAPC of MLGs identified in the three lakes and time periods. Fill colours represent MLGs and correspond to colours in A.

**Table 1.** Genetic diversity of the *Daphnia pulex* population in three lakes of West Greenland integrated over several sediment depths.

| Pop | N | MLG | eMLG | SE |
| --- | --- | --- | --- | --- |
| SS4 | 28 | 3 | 1.7 | 0.7 |
| SS1590 | 10 | 6 | 6.0 | 0.0 |
| SS1381 | 21 | 1 | 1.0 | 0.0 |
| Total | 59 | 10 | 3.9 | 1.1 |

The *Daphnia* lineages of the three lake populations formed well-defined clusters (DAPC, Fig. 3B), separating the lakes at almost equal distances. The analysis also illustrates the genetic similarity of three MLGs of SS4. While the clones of SS1590 were more scattered they still formed their own cluster without overlap to the other lakes.

## Discussion

The results of this study indicated almost equal separation of the three *Daphnia* populations among the lakes (Fig. 3) and despite considerable in-lake and regional environmental change since ~1700 AD (discussed below), the clonal structure of *Daphnia* populations in two of three lakes (SS4 and SS1381) was remarkably stable. Importantly, we found strong dominance of a single, lake-specific asexual clone throughout the past two to three centuries in each of the two lakes.

### Spatial patterns

The analysis of population genetic structure revealed almost equal separation of the three populations (Fig. 3), a spatial pattern that may reflect the primary environmental differences among the lakes as well as the historic connectivity between these lakes: SS4 is oligosaline and thus differs strongly from SS1381 and SS1590. The latter two are quite similar to each other in terms of their water chemistry (Table S1) and typical of many of the freshwater lakes around the head of the fjord [33]. Clearly, ionic composition and inorganic C chemistry is an important driver of genotypic variability as much as it determines regional biodiversity and community structure (diatoms, chrysophytes; [51,52]. Conductivity is the dominant control on algal composition in this area and is highly correlated with both DOC concentration and CDOM quality [37]. Conductivity was also identified as one of the major factors driving clonal composition of *Daphnia* populations in an area about 250 km north of Kangerlussuaq [30]. Lakes SS1381 and SS1590 have never been connected hydrologically, since SS1381 lies outside the boundary of the large palaeolake that included SS4 in the period immediately after deglaciation [41]. The lack of shared clones between lakes suggests local adaptation to the environmental conditions of each resident lake, in particular given the geographic proximity between all three study lakes (less than 10 km) and a likely lack of dispersal limitation as observed in other populations of West Greenland *Daphnia* [29].

The overall greater clonal diversity of the SS1590 *Daphnia* population compared with that of SS4 and SS1381 may reflect the greater diversity of habitats in this lake – its two sub-basins are quite different in terms of their morphometry and substrate variability, including the dominant macrophytes (Anderson, unpublished field observations). Not only are littoral and benthic habitats more diverse in SS1590, but the lake also has a more dynamic response to evaporative driven lake level changes than many of the lakes in the area. These short-term lake level changes also exacerbate habitat heterogeneity more at this site than others, due to its morphometry. SS1381 has a simpler morphometry, essentially with one deep basin. SS4, the largest of the lakes can be considered a pelagic system and while lake levels are variable here as well, the contribution of the littoral zone to whole lake production and diversity is probably more limited.

### Regional environmental change

Lakes in the Kangerlussuaq area are tightly coupled to regional climate change which influences both terrestrial landscape and in-lake aquatic processes, such as hydrological runoff, terrestrial productivity, lake levels, conductivity and DOC concentration [53]. The period covered by the genetic analyses, ~300 years, includes the end of the Little Ice Age (LIA); a period of considerable change in regional climate, with aridity and fluctuating temperatures. Lake levels would have been lower than present [41,43] and aridity affects dust deflation from the sandurs immediately north and to the east of the study area. As the lakes are strongly P-limited [54], dust is a possible important nutrient source. Atmospheric reactive N deposition has increased across the Arctic [55] and Greenland is no exception. The changing nutrient balance associated with these varying sources will have impacted primary production and thus secondary producers (i.e. *Daphnia*). But as well as these broader, regional responses which can be traced across lakes, each individual lake has its own ecological trajectory in environmental space (see for example [35]). Long-term air temperature records show considerable variability during the 20^th^ century, including a period of pronounced cooling. Rapid warming characterizes the start of the 21^st^ century. Lake temperatures track air temperatures tightly in this area [56] and so epilimnetic temperatures would have varied similarly.

As an example for local environmental variability since LIA, SS4 showed pronounced variation of okenone, a pigment which can be used as a proxy for purple sulphur bacteria (PSB) abundance [34]. In SS4, in conjunction with indicators of primary production, variation of ephippial densities provides evidence of historic changes in trophic interactions. Ephippial production serves as a rough estimate of *Daphnia* population size [57,58]. There is some suggestion that over the last ~800 years the *Daphnia* population in SS4 has been tracking purple sulphur bacteria (Frisch & Osburn unpublished), which *Daphnia* may exploit as a direct or indirect food source, as observed in other studies [59,60].

### Long-term persistence of dominant clones

Despite these profound environmental changes, there is limited genotypic variability over time. We did not find evidence for environmental dynamics to drive patterns of population genetic structure or genetic diversity over the past several centuries. Our data suggest the presence of a single clone in two of the study lakes, in SS4 (MLG1) and SS1381 (MLG10), across all sample depths, over the past 200 to 300 years. In SS4, two other clones were detected in two different time periods, both with very low abundance. In contrast, the third lake (SS1590) had a higher number of clones. While it is obvious that clonal diversity in this lake overall was much higher, the temporal pattern of genetic structure and diversity cannot be identified with certainty due to the low numbers of isolates; overall ephippial density in this lake was much lower (personal observation) with a smaller amount of well-preserved eggs suitable for genetic analyses.

There are several tenable explanations for the apparent dominance and persistence of single clones:

#### (1) The persistent clone may be a general-purpose genotype that successfully exploits a variety of historic environments that are not (yet) within a harmful range

The invasion and dominance of a single, asexual *Daphnia* clone has been observed in several lakes throughout a wide geographic range. For example, an asexual *Daphnia pulex* clone invaded the population in Lake Naivasha, Kenia in the 1920s, and then displaced the local sexual, genetically diverse *Daphnia* population [20]. Successful invasion of the same clone was observed throughout Africa in a wide range of aquatic habitats and environmental conditions, testifying to the exceptional niche breadth of this asexual clone. Similar observations have been made in Japan [61], where four asexual *Daphnia* clones invaded a large number of aquatic habitats across a wide geographical range and ecological conditions. In our study of Arctic populations, the persistence of *Daphnia* clones despite major temperature changes over the last centuries in the study area may reflect the considerable in-lake temperature gradient that *Daphnia* experience on a regular basis. At SS4, daily vertical migration to graze directly or indirectly on PSB and POC (DOC) in the metalimnion at SS4 (a distance of some 8-12 m) would expose animals to a temperature change of >10 °C, considerably more than the temperature change during the recent millennia. Moreover, the annual temperature range in the epilimnion is also in the order of 10 °C [44]. Although only SS4 has a metalimnetic PSB plate, all three study lakes have substantial gradients in DOC and POC, suggesting that they are utilising a microbial loop in the lakes. The three study lakes are seasonally anoxic with hypolimnetic reductions in O2 during both summer and winter. It is possible that these seasonal environmental changes and the vertical O2 gradients exert greater physiological stress in *Daphnia* than the stress associated with regional warming. Of course, this genetic stasis may not continue should environmental change in SW Greenland stay at a similar rate [31].

#### (2) Undetected by the applied genetic resolution, a single MLG may comprise several distinct clonal lineages, each adapted to a specific historic environment

The possibility of cryptic genetic variation must be considered due to the limited resolution offered by microsatellite markers which cannot fully account for possibly existing genome-wide variation. For example, using 12 microsatellite loci, So et al. [61] detected only four MLGs in asexual *Daphnia pulex* that had invaded Japanese lakes and ponds. Interestingly, these MLGs comprised 21 mitochondrial haplotypes, indicating a higher genotype diversity than that resolved by the microsatellite loci [61], a finding that was later confirmed using whole genome sequencing [62]. In particular, these authors concluded that the observed divergent traits in these closely related asexual genotypes evolved without recombination from an ancestral clone by a limited number of functionally significant mutations [62].

#### (3) Epigenetic modifications explain clonal adaptation and allow clones to persist

There is increasing evidence that epigenetic mechanisms contribute to evolution [63,64] and to adaptive responses related to climate and environmental change [65–67]. In the absence of genomic variation, it is becoming increasingly evident that changes in epigenetic profiles could allow for rapid, heritable adaptations to environmental cues that precede the more slowly evolving changes in DNA sequences [68], i.e. the classical Darwinian mutation accumulation and selection concept. Epigenetic modifications would allow persistence of asexual clones in the absence of recombination and generation of genetic variation, because transgenerational inheritance of DNA methylation is highly likely in *Daphnia*, whose gametes are derived from almost fully matured tissue [69].

### Synthesis

In order to test the ecological implications of these findings, laboratory experiments are needed in future studies. In particular, representatives of the current populations can be used to test whether the dominant clones exhibit phenotypic plasticity with a wide tolerance towards local conditions (temperature, food, salinity), and whether local adaptation to these conditions can be observed. If possible, hatchlings of eggs from older sediments should be included to compare phenotypic and transcriptomic responses of ancient and contemporary isolates [12,70]. In addition, the role of epigenetic modifications should be considered in further studies.

These results, to our best knowledge, are the first to report the population genetic structure of asexual, polyploid *Daphnia* populations of the circumpolar *Daphnia pulex*-complex over century timescales. Future, more intensive sampling may reveal additional clonal lineages; however, the observed dominance patterns and persistence of individual clonal lineages are unlikely to change. Our results stimulate several questions relating to the mechanisms of adaptation in these populations, as well as their evolutionary fate during the next decades and beyond, on the assumption that climate change in Greenland and other Arctic regions proceeds on the predicted trajectory with severe ecological consequences.

## Acknowledgements

We are grateful to Erika Whiteford, Madeleine Giles, Amanda Burson and Tania Cresswell-Maynard for help with field work, to Chris O’Grady for assistance in the laboratory, to Andy Moss for providing laboratory space for core processing at UoB and to Fengjuan Xiao (Lboro) for the loss-on-ignition analyses. DF received funding from the European Union’s Horizon 2020 research and innovation programme under the Marie Skłodowska-Curie grant agreement No. 658714. NJA’s work in Kangerlussuaq is the supported by NERC.

## Supplementary Information

Fig. S1 Photograph of a sediment core recovered in SS4

Fig. S2. A-B. Correlation of replicate cores at SS4 C. Core correlation at SS1381

Table S1 Nutrient and major ion water chemistry in the three study lakes

Table S2 Details on allelic diversity for eight microsatellite loci of the three *Daphnia pulex* s.l. populations in this study

## Authors’ contributions statement

DF conceived the study. MD conducted the molecular work and analysed the molecular data with DF and input from JKC. NJA and CLO analysed the environmental data. DF and NJA conducted the field work, interpreted the data and wrote the manuscript with input from all authors. All authors gave final approval for publication and agree to be held accountable for the work performed therein.

