## Supplementary Information for "Centennial clonal stability of asexual *Daphnia* in Greenland lakes despite climate variability"

Fig. S1. Photograph of a sediment core recovered in SS4, confirming an intact sediment-water interface and undisturbed sediment layers.

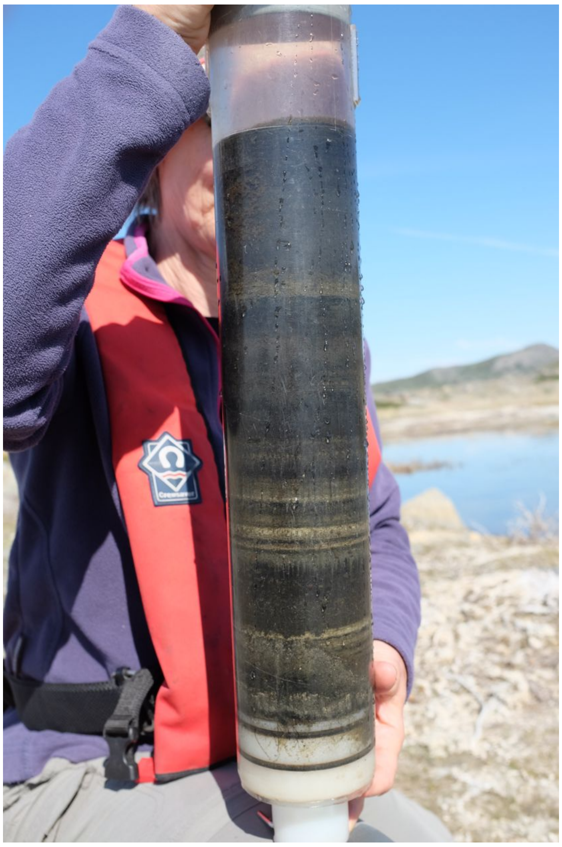

Fig. S2A. Correlation of replicate cores at SS4.

These plots show the clear repeatability of cores taken one location in the basin, approximately within30-m radius. The boat was moved after each core was taken to avoid unnecessary sediment disturbance.

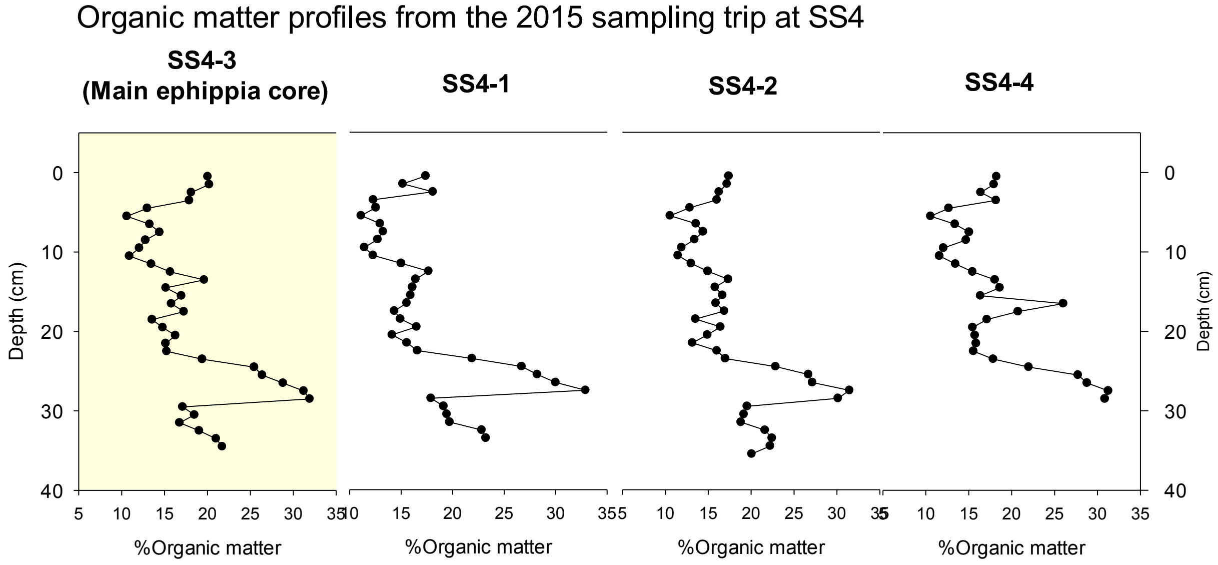

Fig. S2B. Core correlation at SS4.

This figure illustrates the repeatability between cores taken years apart and the relative ease with which organic matter (and/or %C) profiles can be used to transfer an established chronology (in this case a core from the alkenone study and temperature reconstruction by D'Andrea et al. (2011).

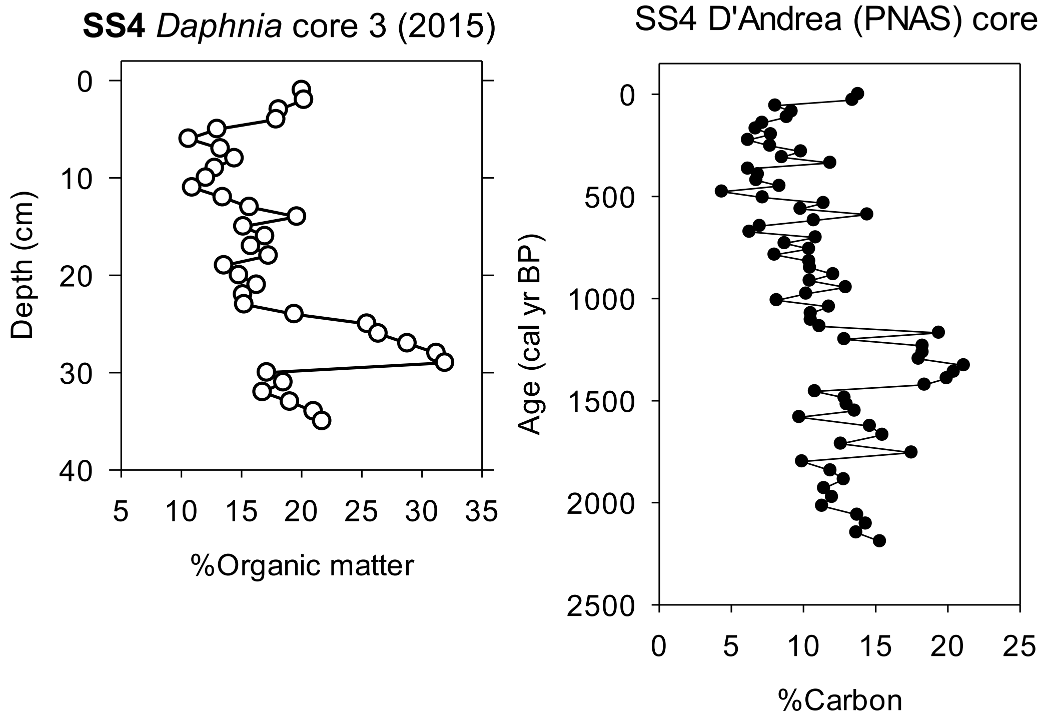

Fig. S2C. Core correlation at SS1381.

As with Figure S2B to illustrate the agreement between the *Daphnia* study core and that from an earlier palaeolimnological study (Law et al. 2015; QSR). The *Daphnia* core has a slower sediment accumulation rate.

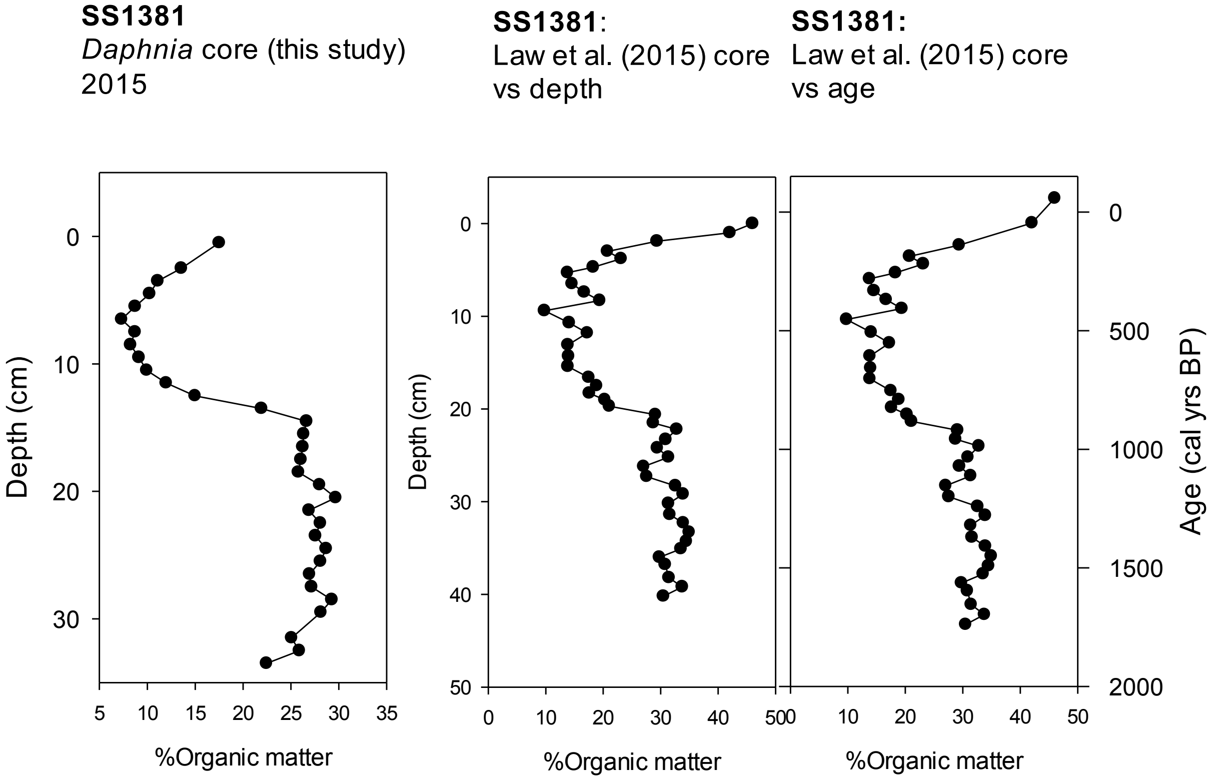

Table S1. Nutrient and major ion water chemistry in the three study lakes SS4, SS1381 and SS1590 near Kangerlussuaq, Southwest Greenland.

| Lake | area | Max depth | SpCond | pH | Alk | TP | TN | $NH_4^+$ | Si | $K^+$ | $SO_4^{2-}$ | DOC |
|---|---|---|---|---|---|---|---|---|---|---|---|---|
| | ha | m | µS/cm | | $\mu eql^{-1}$ | µg/L | $\mu meql^{-1}$ | µg/L | | µg/L | µg/L | mg/l |
| SS4 | 90 | 24.0 | 3210 | 9.04 | 17330 | 10.0 | 1100.0 | n.a. | n.a. | 3007.0 | 1650.0 | 71.0 |
| SS1381 | 21.5 | 18.0 | 523 | 8.25 | 3150 | 2.0 | 947.0 | 3.90 | 0.4 | 554.0 | 31.0 | 28.3 |
| SS1590 | 24.6 | 17.5 | 219 | 7.91 | 1500 | 15.0 | 753.0 | 4.68 | 0.4 | 228.0 | 26.0 | 19.8 |

Table S2. Details on allelic diversity for eight microsatellite loci of the three *Daphnia pulex* s.l. populations in this study. Allele= number of allels per locus, I-D = Simpson index, Hexp = Nei's 1978 gene diversity

| locus | Allele | 1-D | Hexp | Evenness |
|---|---|---|---|---|
| Dp291 | 4 | 0.70 | 0.70 | 0.93 |
| Dp173 | 3 | 0.44 | 0.45 | 0.82 |
| Dp437 | 6 | 0.77 | 0.77 | 0.89 |
| Dp43 | 3 | 0.55 | 0.56 | 0.87 |
| Dp461 | 4 | 0.72 | 0.72 | 0.91 |
| Dp162 | 3 | 0.61 | 0.62 | 0.89 |
| Dp401 | 7 | 0.81 | 0.82 | 0.94 |
| Dp369 | 5 | 0.70 | 0.71 | 0.88 |
| mean | 4.4 | 0.66 | 0.67 | 0.89 |
